# MALT: Fast alignment and analysis of metagenomic DNA sequence data applied to the Tyrolean Iceman

**DOI:** 10.1101/050559

**Authors:** Alexander Herbig, Frank Maixner, Kirsten I. Bos, Albert Zink, Johannes Krause, Daniel H. Huson

**Affiliations:** Institute for Archaeological Sciences, University of Tübingen, Rümelinstrasse 23, 72072 Tübingen, Germany.; Max Planck Institute for the Science of Human History, Kahlaische Strasse 10, 07745 Jena, Germany.; Institute for Mummies and the Iceman, European Academy of Bozen/Bolzano (EURAC), Viale Druso 1, 39100 Bolzano, Italy.; Center for Bioinformatics, University of Tübingen, Sand 14, 72076 Tübingen, Germany

## Abstract

Modern next generation sequencing technologies produce vast amounts of data in the context of large-scale metagenomic studies, in which complex microbial communities can be reconstructed to an unprecedented level of detail. Most prominent examples are human microbiome studies that correlate the bacterial taxonomic profile with specific physiological conditions or diseases.

In order to perform these analyses high-throughput computational tools are needed that are able to process these data within a short time while preserving a high level of sensitivity and specificity.

Here we present MALT (**ME**GAN **AL**ignment **T**ool) a program for the ultrafast alignment and analysis of metagenomic DNA sequencing data. MALT processes hundreds of millions of sequencing reads within only a few hours. In addition to the alignment procedure MALT implements a taxonomic binning algorithm that is able to specifically assign reads to bacterial species. Its tight integration with the interactive metagenomic analysis software MEGAN allows for visualization and further analyses of results.

We demonstrate MALT by its application to the metagenomic analysis of two ancient microbiomes from oral cavity and lung samples of the 5,300-year-old Tyrolean Iceman. Despite the strong environmental background, MALT is able to pick up the weak signal of the original microbiomes and identifies multiple species that are typical representatives of the respective host environment.

## Background

Modern DNA sequencing technologies rapidly produce vast amounts of data, allowing scientists to sequence samples to an unprecedented level of detail. In metagenomics, DNA sequencing is used to study environmental samples, such as soil (Mackelprang, Waldrop et al. 2011), water samples (Sunagawa, Coelho et al. 2015) or host-associated samples, in particular from humans (The Human Microbiome Project Consortium 2012). Taxonomic analysis of such data aims at determining the microbial composition of the given samples. One main question is whether changes in composition are correlated to specific attributes of the samples, such as disease state in human-associated studies.

In metagenomics analyses, it is useful to distinguish between taxonomic profiling and taxonomic binning. The goal of taxonomic profiling is to estimate the microbial content of samples based on the given sequencing reads, but without attempting to assign the set of reads to specific organisms. For example, MetaPhlan2 (JÛnsson, Ginolhac et al. 2013) bases its taxonomic analysis on the alignment of a subset of the reads to a set of specific marker genes, while other tools such as KRAKEN (Wood and Salzberg 2014) analyze the k-mer content of samples. While taxonomic binning also provides an estimation of microbial community structure of a sample, its primary aim is to assign as many reads as possible to specific taxa. This type of analysis is usually based on an alignment of the reads against a database of reference sequences (such GenBank, in the case of DNA alignments, or NCBI-nr, in the case of protein alignments (Pruitt, Tatusova et al. 2007, Benson, Cavanaugh et al. 2013)).

In this study we will apply a taxonomic binning approach to metagenomes of two ancient human specimens to reconstruct the microbial community structure. Metagenomes of ancient samples usually display a mix of low amounts of ancient endogenous DNA (aDNA) and a high background contamination (Prüfer, Stenzel et al. 2010). Contaminating DNA can originate from humans handling the sample or, if ancient bacterial pathogens are the target, from closely related bacteria in the soil. For this reason, methods have been developed to estimate contamination based on aligning sequences to suitable reference sequences, and to validate the authenticity of ancient DNA, usually by detecting characteristic damage patterns caused by the deamination of cytosines (e.g. mapDamage 2.0 (JÛnsson, Ginolhac et al. 2013)). Additional methods use the evenness of coverage of aligned reads across the reference genome or length distributions for aligned reads, as aDNA tends to be more fragmented than contemporary DNA (Prüfer, Stenzel etal. 2010).

Metagenomic dataset normally are comprised of millions of DNA sequencing reads. Hence, taxonomic binning based on alignment is computationally challenging. The gold standard tool for such analysis is BLAST (Altschul, Gish et al. 1990), due to its sensitivity and useful statistical model. However, the computational time required by BLAST to analyse metagenomic data is prohibitive.

Here we present MALT (**ME**GAN **AL**ignment **T**ool), a program for the fast alignment and analysis of metagenomic DNA sequencing data. MALT contains the same taxonomic binning algorithm (the naïve LCA (Lowest Common Ancestor) algorithm) that is implemented in the interactive metagenomics analysis software MEGAN (Huson, Beier et al. 2016). When aligning against a protein reference database, MALT can also perform functional analysis using a classification such as SEED (Overbeek, Begley et al. 2005). The MALT output of the aligned reads can be further visualized and analyzed in MEGAN.

Like BLAST, MALT computes “local” alignments between highly conserved segments of reads and references. However, MALT can also calculate “semi-global” alignments in which reads are aligned end-to-end. Semi-global alignments are more suitable for assessing various quality and authenticity criteria that are commonly applied in the field of paleogenetics. Semi-global alignments are also useful when aligning 16S rRNA data against a reference database such as SILVA (Quast, Pruesse et al. 2013). All alignments can be output in SAM format (Li, Handsaker et al. 2009).

MALT is written in Java and installers are available for Linux, MacOS and Windows.

MALT is freely available from: http://ab.inf.uni-tuebingen.de/software/malt

Another recent tool that performs fast alignments of DNA reads to a nucleotide target database is *lambda* (Hauswedell, Singer et al. 2014). Since the program reports only local alignments and not semi-global ones the applicability of *lambda* to aDNA is limited. Moreover, *lambda* does not perform taxonomic assignment.

In this paper, we use MALT to perform a comparative analysis of the metagenomes of two tissue samples from the 5,300-year-old Tyrolean Iceman, a European Copper Age ice mummy (Spindler 1994). To assess the performance of MALT the same datasets are analyzed with BLAST and the fast DNA sequence aligner *lambda*.

## Algorithm

MALT is based on the seed-and-extend paradigm, as made popular by BLAST. MALT consists of two programs, *malt-build* and *malt-run*.

First, *malt-build* is used to construct an index for the given database of reference sequences (which is input in FastA format). To do so, *malt-build* determines all occurrences of spaced seeds (Burkhardt and Kärkkäinen 2001, Ma, Tromp et al. 2002, Buchfink, Xie et al. 2015) in the reference sequences and places them into a hash table (Ning, Cox et al. 2001), keeping only seeds that do not occur too often (1000 by default).

*malt-run* is then used to align a set of query sequences against the reference database. To this end, the program generates a list of spaced seeds for each query and then looks them up in the reference hash table, which is kept in main memory. Using the x-drop extension heuristic (Altschul, Gish et al. 1990), a high-scoring ungapped alignment anchored at the seed is computed and is used to decide whether to extend to a full alignment. Local or semi-global alignments are computed using a banded implementation (Chao, Pearson et al. 1992) of the Smith-Waterman (Smith and Waterman 1981) or Needleman-Wunsch (Needleman and Wunsch 1970) algorithms, respectively. The program then computes the bit-score and expected value (E-value) of the alignment and decides whether to keep or discard the alignment depending on user-specified thresholds for bit-score, E-value or percent identity. The application of *malt-run* is illustrated in Figure 1.

**Figure 1.**
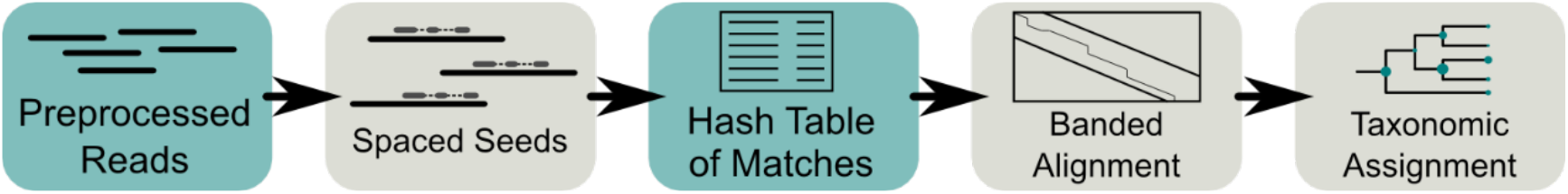
**Schematic overview of MALT**. For each preprocessed metagenomic sequencing read, the algorithm generates all contained spaced seeds and looks them up in a hash table of spaced seeds representing the reference database. A banded alignment is calculated for each match of seeds. Once all alignments for a given read have been calculated taxonomic binning of the read is performed using the LCA algorithm.

In order to use MALT in aDNA studies to screen for bacterial DNA and to assess the taxonomic composition of ancient microbial communities we apply the following workflow (Figure 2). First we use *malt-build* to construct a MALT index on all complete bacterial genomes in GenBank (Benson, Cavanaugh et al. 2013). This has to be done once and we rebuild the index only for updating the target database. For each sample we preprocess all reads by trimming sequencing adapters and merging overlapping paired-end reads. Then we align the resulting sequences against the reference database using *malt-run* in semi-global mode. MALT generates output in RMA format and in SAM format. The former can be used for interactive analysis of taxonomic composition in MEGAN and the latter for alignment-based assessment of damage patterns etc.

**Figure 2.**
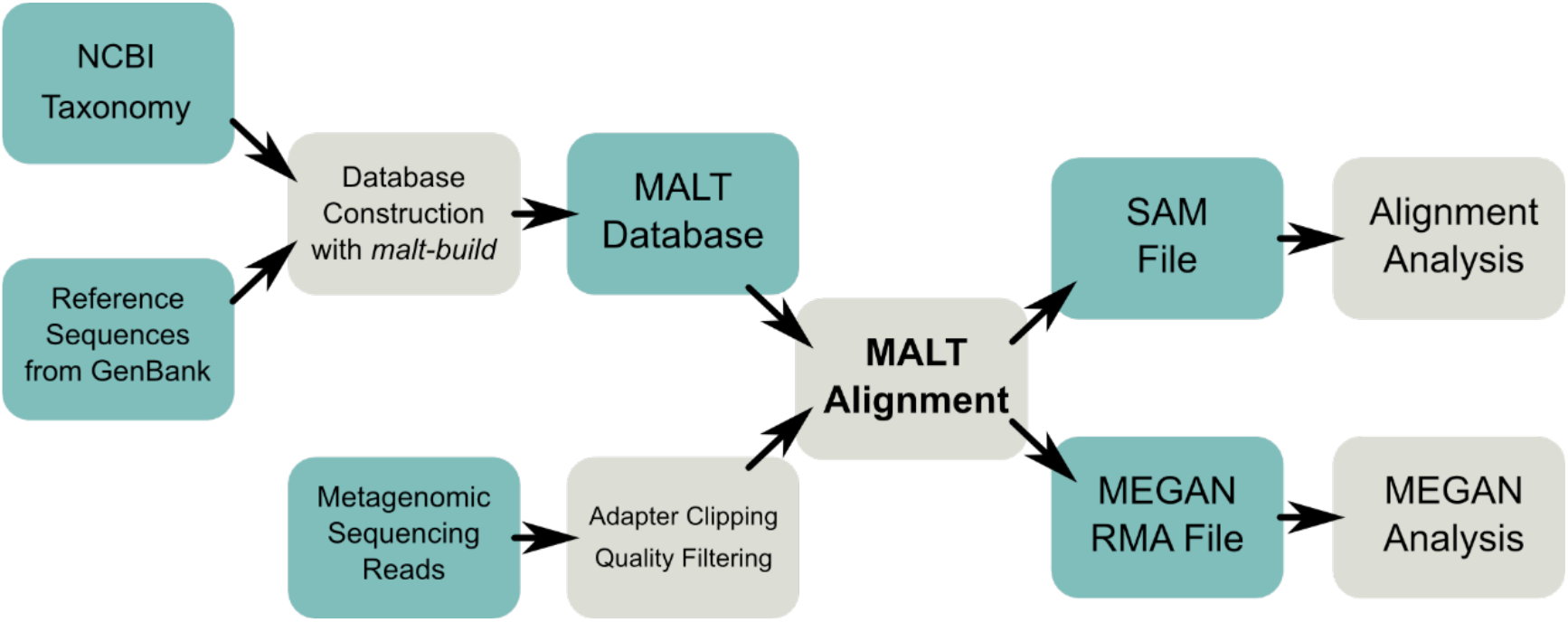
**Schematic overview of our MALT-based analysis workflow**. A reference index is generated for all bacterial genomes from GenBank. Preprocessed metagenomic sequencing reads are aligned against the reference sequences. An RMA result file is produced for further analysis in MEGAN. Alignments are also stored in SAM format.

## Results

The datasets from the lung and gingiva tissue of the Tyrolean Iceman used as input for MALT comprise 8 and 12 million metagenomic DNA sequencing reads, respectively. Based on random subsamples of reads, we compared the alignment speed of MALT, BLAST and *lambda* (Table 1). In our test environment MALT processed 10 million reads in about 20 minutes. This is about 25 times faster than *lambda* and more than 120 times faster than BLAST. While the runtime of BLAST grows almost linearly with the number of reads, MALT benefits from dynamically loading the index into main memory, which increases the runtime performance asymptotically. A similar effect can be observed for *lambda*. For our target database MALT requires about 250 GB of RAM.

**Table 1.** Runtime comparison of BLAST, *lambda* and MALT on test datasets of different size. Runtimes are given in seconds.

| number of<br>reads | runtime<br>BLAST | runtime<br>lambda | runtime<br>MALT |
| --- | --- | --- | --- |
| 100.000 | 1.729 | 1.024 | 38 |
| 1.000.000 | 16.077 | 6.414 | 155 |
| 10.000.000 | 150.045 | 31.163 | 1.252 |

In order to assess the sensitivity of the two methods we compared their results to the reads aligned by BLAST. Considering only alignments with an e-value of 10-5 or smaller, MALT finds at least one alignment in 82% of the cases for which BLAST finds such an alignment, while the corresponding rate for lambda is less than 20%.

The output of MALT was analyzed using the metagenomic analysis software MEGAN. In both samples 90% of the reads could not be assigned to any taxa and were therefore not considered during further analysis (Table 2). Here it has to be considered that our database only consists of bacterial genomes and excludes any eukaryotes and viruses. For both samples, the studied microbiomes are dominated by environmental bacteria (Figure 3) (Hyde, Haarmann et al. 2013, Metcalf, Xu et al. 2015). About 55% of the reads for the oral cavity sample and about 95% of the reads for the lung tissue sample were assigned to species of the genus *Pseudomonas*. About 15-30% of the remaining reads were assigned to *Clostridia*. In the oral cavity sample a substantial amount of reads was assigned to *Carnobacteria* typically involved in body degradation, which represented only a minor fraction in the lung tissue.

**Table 2.** Number of input reads and number of taxonomically assigned reads for the oral cavity and lung dataset. In addition, the number of assigned reads for various selected genera and species is provided. A full list of taxonomic read counts is provided in Table S1.

|  | **Oral Cavity** | **Lung** |
| --- | --- | --- |
| Total Reads | 12,486,937 | 8,503,240 |
| Assigned Reads | 1,098,791 | 1,867,705 |
| *Streptococcus* | 5,551 | 55 |
| *Streptococcus mutans* | 159 | 0 |
| *Staphylococcus* | 4,109 | 27 |
| *Staphylococcus aureus* | 341 | 0 |
| *Treponema denticola* | 78 | 0 |
| *Filifactor alocis* | 561 | 0 |
| *Fusobacterium nucleatum* | 1,941 | 0 |
| *Lactococcus lactis* | 843 | 0 |
| *Haemophilus* | 224 | 66 |
| *Klebsiella pneumoniae* | 172 | 375 |

**Figure 3.**
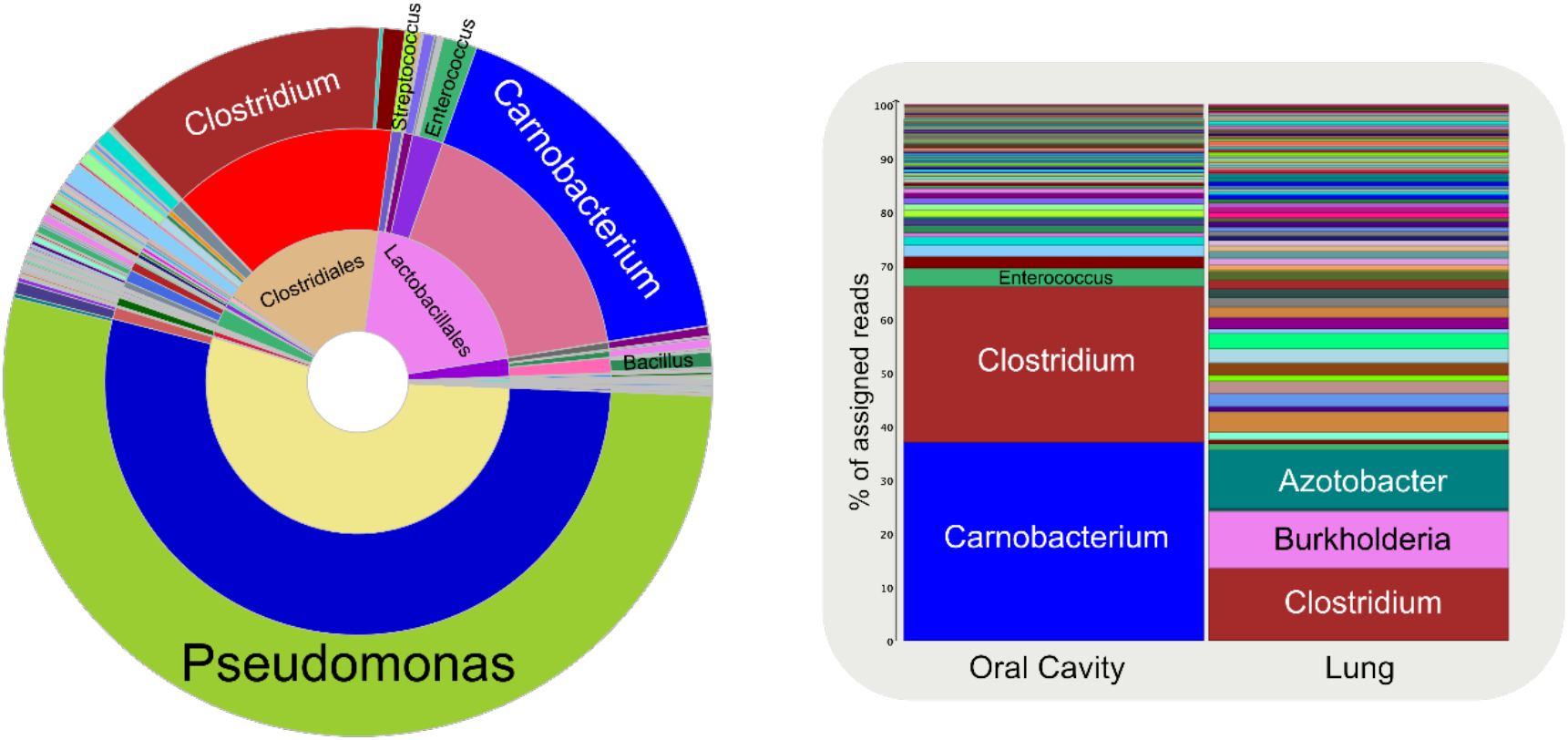
**Taxonomic analysis of oral cavity and lung tissues from the Iceman. Left:** Taxonomic composition of the oral cavity sample. About 55% of the assigned reads represent species of the *Pseudomonas* genus. Further genera with a substantial representation are *Clostridium* and *Carnobacterium*. **Right:** Taxonomic composition of oral cavity and lung samples after omission of reads assigned to Pseudomonas. *Clostridia* represent a substantial part of the microbiome in the lung sample. Further genera found in larger amounts are *Azotobacter* and *Burkholderia*.

MALT is sensitive enough to detect weak signals of original microbial community members of the studied tissues (Table 2). Various *Streptococcus* species, e.g. *Streptococcus mutans*, were detected in the oral cavity sample. Reads were also assigned to *Staphylococcus* species, such as *Staphylococcus aureus*. In addition, there is a weak signal for *Treponema denticola*. We also have read assignments for *Filifactor alocis, Fusobacterium nucleatum* and *Lactococcus lactis*. The presence of these bacterial species is typical for the human oral microbiome (Dewhirst, Chen et al. 2010).

In the lung tissue we see only very weak signals for *Streptococci* and *Staphylococci* and no assignments for the species detected in the gingiva sample. However, in the oral cavity as well as the lung sample we detect a weak signal for the genus *Haemophilus* with a few specific assignments to the species *Haemophilus influenzae*. This Gram-negative bacterium is found in the human respiratory system and represents a relevant pathogen, which can cause bacterial meningitis (Erwin and Smith 2007).

We see a stronger signal in the lung tissue compared to the oral cavity for *Klebsiella pneumoniae*. This Gram-negative non-motile bacterium is part of the human oral flora and also of the skin and gut microbiome. It is usually nonpathogenic, but when it populates the lung, it can cause destructive changes to the alveoli and represents a common cause of pneumonia (Podschun and Ullmann 1998).

We applied mapDamage 2.0 (JÛnsson, Ginolhac et al. 2013) to the MALT alignments for the aforementioned species and detected characteristic damage patterns for all of them (Figure S1). Compared to the human DNA (Figure S2) the damage signals are slightly lower, which might be due to cross-mapping of DNA fragments from closely related modern species.

## Discussion

Metagenomic analysis is facing a wide range of challenges, in particular, the comparison of huge amounts of sequencing data against a steadily increasing number of reference sequences. While the overall composition can be characterized without precise alignments, other types of analysis require the assessment of complete alignments of the metagenomic sequencing reads against a comprehensive reference database. Speeding up and streamlining the alignment and analysis of reads will help address the bottleneck of data processing in studies that deal with metagenomic data.

Here, we introduce MALT, a fast DNA read aligner that implements semi-global alignment and can perform taxonomic binning in parallel. We compared MALT to the gold standard BLASTN and also to the recently developed DNA sequence alignment software *lambda*, which is also intended for use on large datasets. MALT is substantially faster than *lambda* and achieves a runtime allowing for the processing of a complete HiSeq run within a few days. MALT performs the taxonomic binning simultaneously, while this step is not included by *lambda* and would thus require an additional step and more time.

In the field of paleogenetics, accurate alignments are required to determine characteristic damage patterns of ancient DNA (aDNA) in order to assess the authenticity of the detected fragments. In addition, large amounts of data are produced as the proportion of endogenous DNA is usually low and the samples composition is dominated by environmental sources. Data processing pipelines specialized on aDNA data have to deal with these challenges. We recently developed EAGER (Peltzer, Jäger et al. 2016), an efficient and user-friendly ancient genome reconstruction workflow that implements the data preprocessing steps that are required for MALT. PALEOMIX (Schubert, Ermini et al. 2014) is another aDNA pipeline, which focuses more on the metagenomic aspects of data analysis. MALT can be easily integrated into such pipelines.

We demonstrated in this study that MALT is able to characterize the overall composition of a metagenomic sample and is sensitive enough to pick up weak traces of ancient microbiomes that are otherwise hidden due to a strong environmental background. For this, MALT was applied to metagenomic datasets of a lung and an oral cavity sample of the Tyrolean Iceman. We see evidence of multiple genera and species that are commonly found in the human oral flora or that are potentially able to cause disease. However, those detections have to be seen as mere candidates.

The low coverage of the target genomes does not allow for a precise genomic characterization of the bacteria. Furthermore, in some cases it is impossible to distinguish between background and target microbiome. For example, *Klebsiella pneumonia* is a bacterium that is not only part of various human microbiota but it is also able to survive in soil (Podschun and Ullmann 1998).

On the other hand, the presence of the bacterium *Treponema denticola* in the Icemans oral cavity has already been shown in a previous publication (Maixner, Thomma et al. 2014). This study presented a partial reconstruction of the *Treponema* genome and provided an assessment of the overall bacterial community composition similar to that of our present study.

In order to verify the presence of candidates that have been detected by a screening approach as described above, the genomes of the candidate species have to be reconstructed, at least partially. This would allow for further genome evolutionary analysis as recently shown for the *Helicobacter pylori* strain from the Iceman’s stomach (Maixner, Krause-Kyora etal. 2016).

MALT can make use of various different target databases. Here we used full bacterial genomes, which provide a rather low resolution compared to 16S rRNA databases such as SILVA (Quast, Pruesse et al. 2013). While our database contains 2.782 bacterial genomes, SILVA has more than 4 million bacterial entries (SILVA 126). However, 16S rRNA amplicon sequencing suffers from biases in particular in the context of ancient microbiome studies (Ziesemer, Mann et al. 2015). The extraction of 16S rRNA reads from shotgun data on the other hand restricts the analysis to high abundance taxa. By using full genomes we can also detect low abundance taxa while at the same time allowing for the assessment of authentication criteria through precise alignments.

MALT can be applied in aDNA screening studies such as demonstrated here, but also to pathogen screening in clinical contexts or in large-scale metagenomic and metatranscriptomic projects. All parameters from the seeding and alignment steps to the taxonomic binning algorithm are customizable and allow for a precise adaptation to the individual needs of each study. The tight integration of MALT with the metagenomic analysis software MEGAN allows for easy visualization of results and for additional interactive analyses.

With MALT we provide a powerful method for high-throughput metagenomic data processing, which - in combination with MEGAN - represents a versatile framework that can be easily integrated into existing or newly developed workflows.

## Methods

### Reference database construction

For the reconstruction of the reference database all complete bacterial genomes and plasmids were downloaded from the NCBI FTP server on the 10th of December 2015.

The following entries were excluded:

multiisoloate_uid216090

multispecies_uid212977

uncultured_Sulfuricurvum_RIFRC_1_uid193658

uncultured_Termite_group_1_bacterium_phylotype_Rs_D17_uid59059

*malt-build* (version 0.1.2) was used to create the MALT target database using standard parameters. In addition, the same data were used to create indexed target databases for BLAST (Altschul, Gish et al. 1990) (version 2.2.31+) and *lambda* (Hauswedell, Singer et al. 2014) (version 0.9.2).

### Runtime comparison

For the generation of different test sets 100,000,1000,000 and 10,000,000 reads were sampled from the Iceman oral cavity dataset. The alignment programs of MALT (version 0.1.2), BLAST (Altschul, Gish et al. 1990) (version 2.2.31+) and *lambda* (Hauswedell, Singer et al. 2014) (version 0.9.2) were applied to the different datasets with comparable parameter settings. For all three programs the DNA alignment mode (blastn) was chosen. The maximal E-value was set to 1.0. The maximal number of alignments for each query was set to 100. The minimal percent identity was set to 85. The number of threads was set to 32. The alignment type of MALT was set to Local in order to be comparable to the other programs.

The computations were performed on a Dell PowerEdge R630 with two Intel Xeon E5-2698 v3 2.3 GHz CPUs and 384 GB RAM.

### Sample preparation and DNA sequencing for oral cavity and lung tissues from the Iceman

Biopsy samples of the Icemans gingiva and lung tissue were subjected to DNA extraction at the ancient DNA Laboratory of the EURAC - Institute for Mummies and the Iceman, Bolzano, Italy. Sample preparation and DNA extraction was performed in a dedicated pre-PCR area following the strict procedures required for studies of ancient DNA: use of protective clothing, UV-light exposure of the equipment and bleach sterilization of surfaces, use of PCR workstations and filtered pipette tips. DNA extraction was performed with approximately 40 mg of gingiva and lung tissue and using a chloroform-based DNA extraction method according to the protocol of Tang et al. (Tang, Zeng et al. 2008). Extracts (including one negative control per tissue sample) were converted into double indexed libraries following methods described elsewhere (Meyer and Kircher 2010), using 10μl of template and minor modifications (Bos, Harkins et al. 2014). Libraries were subsequently amplified using *AccuPrime Pfx* (Invitrogen) and sequenced on an Illumina HiSeq 1500.

### Metagenomic analysis of oral cavity and lung tissues from the Iceman

For the analysis of the two metagenomic datasets MALT (version 0.1.2) was applied with the same parameters as for the runtime comparison with the exception that the alignment type was set to SemiGlobal. For the taxonomic assignment the *top percent* parameter was set to 1.0. The *min support percent* parameter was set to 0.01 for the oral cavity sample. As the bacterial composition of the lung tissue sample was dominated by Pseudomonas species (~95%) a less strict *min support percent* setting of 0.001 was chosen for this sample.

Further analysis and visualization of the bacterial composition were performed with MEGAN (Huson, Beier et al. 2016).

## Author contributions

A.H., F.M., K.B., A.Z., J.K. and D.H. conceived of the study. F.M., K.B., A.Z. and J.K. designed experiments. F.M. and A.Z. performed sampling. F.M. and K.B. performed experiments. A.H., F.M., K.B., A.Z., J.K. and D.H. analysed data. D.H. implemented the MALT program. A.H., J.K. and D.H. designed and set up the MALT aDNA analysis pipeline. A.H., J.K. and D.H. wrote the manuscript with contributions from all coauthors.

## Acknowledgements

We acknowledge the following funding sources: the European Research Council (ERC) starting grant APGREID (J.K., A.H., and K.B.); the South Tyrolian grant legge 14 (F.M.; A.Z.)

---

**Figure S1.**
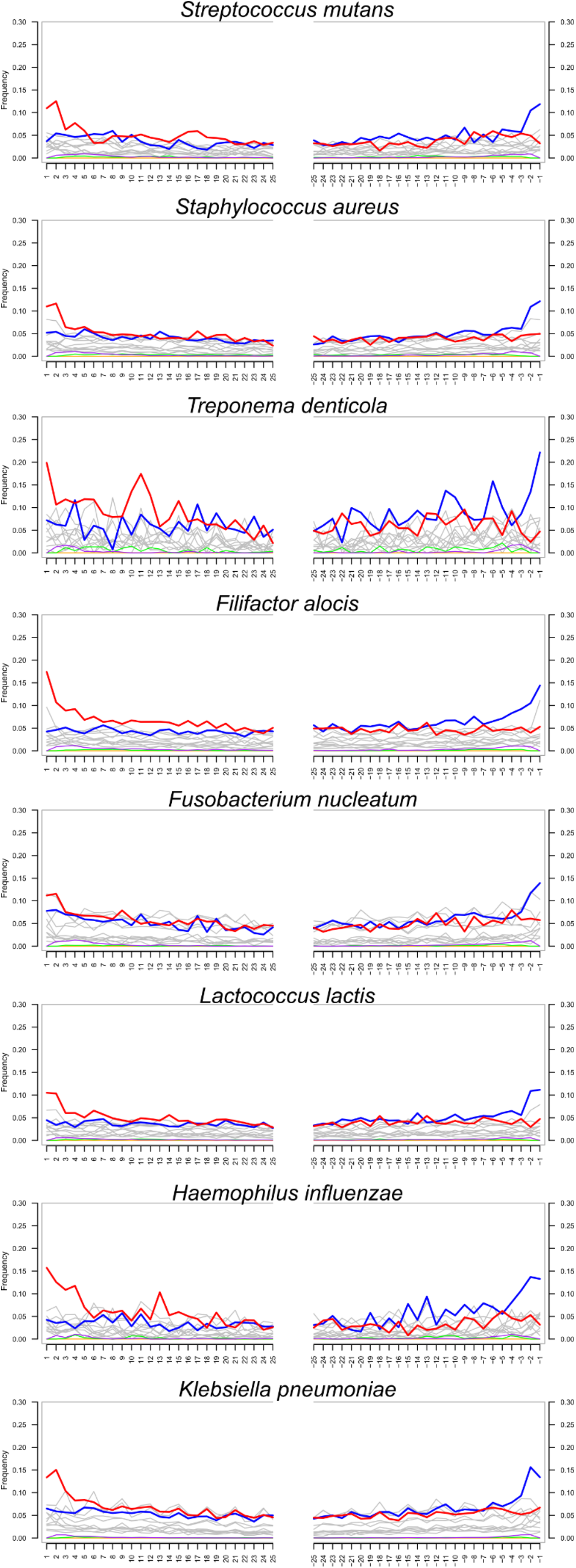
**DNA damage plots generated for various candidate species detected in the Iceman gingiva and lung microbiomes**. The plot for *Klebsiella pneumoniae* was generated for the lung sample, the other plots were generated for the gingiva sample. For each species the plot on the left shows the damage for the 5 end and the plot on the right shows the damage for the 3 end. The x-axis shows the distance to the end of the fragment in nucleotides. The y-axis shows the amount of damage (red: C->T; blue: G->A). For aDNA these two substitutions characteristically occur more frequently at the end of the fragments.

**Figure S2.**
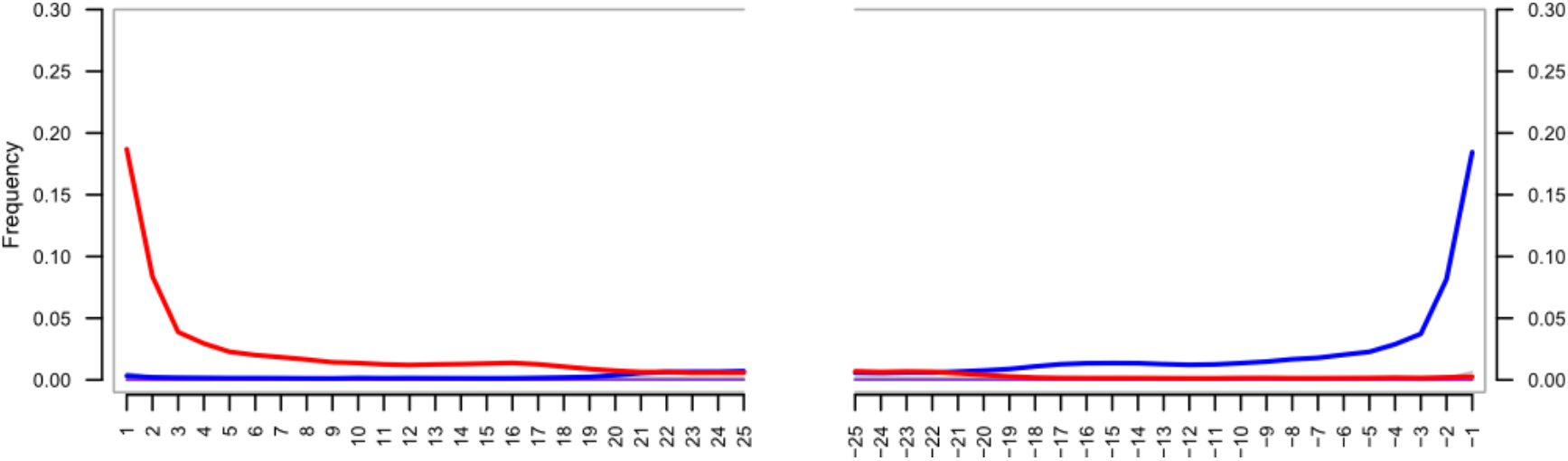
**DNA damage plot generated for the Iceman gingiva reads mapped to the human genome**. The plot on the left shows the damage for the 5 end and the plot on the right shows the damage for the 3 end. The x-axis shows the distance to the end of the fragment in nucleotides. The y-axis shows the amount of damage (red: C->T; blue: G->A). For aDNA these two substitutions characteristically occur more frequently at the end of the fragments.

**Table S1** - Full list of taxonomic read counts for the oral cavity and lung dataset.

